# Gating by induced α-γ asynchrony in selective attention

**DOI:** 10.1101/229526

**Authors:** David Pascucci, Alexis Hervais-Adelman, Christoph M.Michel, Gijs Plomp

## Abstract

Visual selective attention operates through top-down mechanisms of signal enhancement and suppression, mediated by α-band oscillations. The effects of such top-down signals on local processing in primary visual cortex (V1) remain poorly understood. In the present work, we characterize the interplay between large-scale interactions and local activity changes in V1 that orchestrates selective attention, using Granger-causality and phase-amplitude coupling (PAC) analysis of EEG source signals. The task required participants to either attend to or ignore oriented gratings. Results from time-varying, directed connectivity analysis revealed frequency specific effects of attentional selection: bottom-up γ-band influences from visual areas increased rapidly in response to attended stimuli while distributed top-down α-band influences originated from parietal cortex in response to ignored stimuli. Importantly, the results revealed a critical interplay between top-down parietal signals and α-γ PAC in visual areas. Parietal α-band influences disrupted the α-γ coupling in visual cortex, which in turn reduced the amount of γ-band outflow from visual areas. Our results are a first demonstration of how directed interactions affect cross-frequency coupling in downstream areas depending on task demands. These findings suggest that parietal cortex realizes selective attention by disrupting cross-frequency coupling at target regions, which prevents them from propagating task-irrelevant information.

**Significance statement:** In the present work, we demonstrated how selective attention emerges from the interplay between large-scale brain interactions and local structures of information processing in sensory areas. When visual input is relevant, the visual cortex rapidly propagates attended information through feedforward oscillations in the gamma band. When stimuli are irrelevant, however, the parietal cortex suppresses information processing through inhibitory influences in the alpha band. Importantly, we show that alpha activity from parietal cortex disrupts the coupling between low and high frequencies in visual regions, which in turn, determines their amount of feedforward propagation. Our results provide novel insight into how the brain orchestrates selective attention and reveal how the parietal cortex prevents the processing of irrelevant information in other cortical areas.

## Introduction

Our visual environment typically contains more information than our perceptual system can handle. Selective attention is therefore a key mechanism to regulate cortical information flow and prioritize the processing of behaviourally relevant stimuli. How the brain accomplishes such selectivity is one of the fundamental questions in cognitive neuroscience.

At the neuronal level, selective attention operates through the enhancement of activity that represents attended information and the suppression of activity for unattended stimuli. Attentional selection, for example, can either increase or attenuate local responses in early sensory areas, depending on whether they convey information that is relevant or irrelevant for the task at hand (Carrasco, 2011; Daffner et al., 2012; Kanwisher & Wojciulik, 2000; Kastner & Ungerleider, 2001; O’Craven, Downing, & Kanwisher, 1999; Smith, Singh, & Greenlee, 2000). A hallmark of such selective mechanism is the differential pattern of event-related potentials that can be observed on the scalp when a physically identical stimulus is attended or ignored: Attended stimuli often increase the amplitude of early evoked responses (e.g., the P1 component) (Hillyard, Vogel, & Luck, 1998) whereas ignored stimuli evoke a later negative activation, the selection negativity (SN) component(Daffner et al., 2012; Hillyard & Anllo-Vento, 1998), which underlies an inhibitory response to irrelevant and potentially distracting input.

While the modulatory effects of selective attention on sensory analysis and psychophysical performance have been extensively characterized (Carrasco, 2006, 2011; Desimone & Duncan, 1995; Downing, Liu, & Kanwisher, 2001; Driver, 2001; Maunsell & Treue, 2006; Pascucci & Turatto, 2015; Reynolds & Chelazzi, 2004), a fundamental question remains how selective attention is orchestrated between brain areas and what sources and dynamics underlie the emergence of local attentional modulations.

Neuroimaging and lesion studies have contributed to the identification of two distinct functional networks where modulatory signals of selective attention may originate: the ventral attention network (VAN), which comprises the temporo-parietal junction and the ventral frontal cortex and is activated by salient and unexpected stimuli, and the dorsal attention network (DAN), which includes the intraparietal sulcus, superior parietal lobule and frontal eye fields and is engaged by the voluntary and top-down control of attention (Corbetta & Shulman, 2002; Ptak & Schnider, 2011; Vossel, Geng, & Fink, 2014). Recent models of attention suggest that when these attentional control systems are activated, their constituent units may use long-range connections to influence neuronal activity in early sensory areas (Fries, 2009; Vossel et al., 2014).

One way neurons in the attentional network can modulate activity in sensory areas is through coupled oscillations at specific frequencies (Lakatos, Karmos, Mehta, Ulbert, & Schroeder, 2008; Womelsdorf & Fries, 2008). Neuronal oscillations reflect rhythmic synchronization among neuronal ensembles over a wide range of spatial and temporal scales, which may play a crucial role in determining the quality and propagation of sensory signals (Buffalo, Fries, Landman, Buschman, & Desimone, 2011; Foxe & Snyder, 2011; Fries, 2009; Michalareas et al., 2016; Womelsdorf & Fries, 2008). Gamma-band activity (γ, 30-150 Hz), for instance, has been shown to reflect neuronal processing (Gruber, Müller, Keil, & Elbert, 1999; Gruber et al., 1999; Jensen, Kaiser, & Lachaux, 2007; Keil, Müller, Ray, Gruber, & Elbert, 1999) and feedforward communication (Bastos et al., 2015; Michalareas et al., 2016), and has been related to perceptual operations (Singer, 1999) and to neuronal states of high excitability (Fries, Reynolds, Rorie, & Desimone, 2001; Tallon-Baudry, Bertrand, Hénaff, Isnard, & Fischer, 2004; Vossel et al., 2014). Oscillatory activity at lower frequencies (alpha (α) band, 8-14 Hz), in turn, has been implicated in sensory suppression and temporal parsing mechanisms (Haegens et al., 2015; Jensen & Mazaheri, 2010), and has been associated with feedback interactions (Haegens et al., 2015), attentional disengagement (Vanni, Revonsuo, & Hari, 1997) and low states of perceptual receptivity and psychophysical performance (Hanslmayr et al., 2007; Mathewson, Gratton, Fabiani, Beck, & Ro, 2009; Van Dijk, Schoffelen, Oostenveld, & Jensen, 2008). Interestingly, when attention is engaged by relevant stimuli, both a sustained increase in γ power and a decrease in α power have been reported (Bauer, Stenner, Friston, & Dolan, 2014; Foxe, Simpson, & Ahlfors, 1998; Klimesch, Doppelmayr, Russegger, Pachinger, & Schwaiger, 1998; Rajagovindan & Ding, 2011; Thut, Nietzel, Brandt, & Pascual-Leone, 2006; Wyart & Tallon-Baudry, 2008).

These functional roles of α and γ activity suggest neuronal oscillations as the candidate mechanisms through which selective attention operates: sources of attentional control may drive enhancing or suppressive signals at specific frequencies, which in turn could interact with rhythmic synchronization and neuronal communication structures in the ascending pathway (Fries, 2005; Siegel, Donner, Oostenveld, Fries, & Engel, 2008; Womelsdorf & Fries, 2008). However, a good understanding of how directed cortical interactions dynamically implement selective attention is currently lacking.

In the present work, we investigated the brain dynamics of directed interactions that characterize the emergence of selective attention. We recorded functional magnetic resonance imaging (fMRI) and high-density EEG separately, while participants either attended to or ignored physically identical stimuli. Stimulus-evoked EEG source activity was extracted from twenty functionally defined nodes in the attentional and perceptual networks.

Dynamic connectivity analysis revealed a rapid emergence of selective directed connections in the γ- and α-band: Feed-forward interactions in the γ-band increased in response to attended stimuli and were directed from early visual areas to the lateral occipital and fronto-parietal cortex; Inhibitory interactions in the α-band dominated network activity in response to irrelevant stimuli and were orchestrated by parietal cortex. Interestingly, long-range α-band interactions from parietal cortex disrupted phase-amplitude coupling (PAC) between α and γ activity in visual areas, a key mechanism regulating information transmission and neuronal communication (Bonnefond & Jensen, 2015; Dvorak & Fenton, 2014a; Jensen & Mazaheri, 2010). Such reduced local αγ-band asynchrony, in turn led to a suppression of γ-band interactions from visual areas, effectively inhibiting feed-forward information flow.

## Materials & Methods

### Participants

Sixteen healthy subjects (mean age = 31± 20, 3 female), all right-handed and with good visual acuity (mean 1.5 ± range 0.8-1.7, as measured with the Freiburg acuity test (Bach, 1996)) took part in the experiment for monetary compensation. Written informed consent was obtained from each participant before the experiment. The study was performed according to the declaration of Helsinki and after approval by the ethics committee of the University of Geneva.

### Experimental design

Stimuli were Gaussian-windowed sinusoidal gratings (Gabors; σ = 3°, frequency = 3 cycles per degree) presented for 200 ms around a central fixation spot (0.2°). Each Gabor had a contrast of 100% and a variable orientation (maximal -45° to 45° off vertical). There were two conditions depending on whether the Gabor stimulus was relevant or not for the current task. In the Relevant condition, participants had to report the orientation of each Gabor in a two-choice task (left vs. right) by pressing the corresponding key on a response box. In the Irrelevant condition, participants had to detect a color change in the fixation spot (from black to red) occurring on ~33% of trials during the inter-stimulus interval between Gabors.

The same task and stimuli were used in separate fMRI and EEG sessions. Stimuli were back-projected onto a screen (60 Hz, 1024x768 pixels) during fMRI recordings, and presented on a CRT monitor (75 Hz, 1600x1200 pixels) during the EEG session. Stimulus generation, presentation and timing were controlled with PsychoPy (Peirce, 2008) software run under Python 2.7.

The fMRI session consisted of 16 blocks (8 for each condition) of variable length (between 14.7 and 25.2 seconds), presented in pseudo-random order and interleaved with 8 blocks of rest (between 12.6 and 21 seconds). In the EEG session the two conditions were performed in 4, pseudo-randomly interleaved blocks of 100 trials. Written task instructions were provided before each block.

The orientation of Gabors in the Relevant condition was determined through an adaptive staircase procedure (Watson & Pelli, 1983) designed to keep participants’ accuracy at 83%. In the Irrelevant condition, Gabors were randomly tilted toward the left or right with the offset fixed at threshold estimated in the last staircase procedure.

### fMRI data acquisition and preprocessing

fMRI images were acquired using a whole-body Tim Trio system (3T; Siemens Healthcare) at the Brain and Behaviour Laboratory at the University of Geneva, with a radio-frequency (RF) body transmitter and a 32-channel receiver head coil. Functional runs consisted of 295 volumes with 36 T2*weighted echo planar slices (EPIs; repetition time (TR) = 2100 ms; time to echo (TE) = 30 ms; flip angle (FA) = 80°; 3.2 mm slice-thickness with a 0.6 mm gap). After the experimental session, a structural whole-head image was acquired for each participant (TR = 1900 ms; TE = 2.27 ms; field of view (FOV) = 256 x 256 mm^2^; inversion time (TI) = 900 ms; voxel size = 1 x 1 x 1 mm^3^; sagittal orientation).

Functional and structural images were analyzed with the Statistical Parametric Mapping toolbox (SPM12; University College of London, London, United Kingdom). All EPI volumes were realigned to the mean functional image using a two-pass procedure to correct for movement artefacts. The mean of the realigned volumes was then co-registered to the structural image. All images were normalized relative to the standard Montreal Neurological Institute (MNI) space using trilinear interpolation and smoothed with an isotropic 8 mm^2^ full-width half-maximum Gaussian kernel. The time course of cerebrospinal fluid (CSF) and white matter (WM) signals were extracted for each participant before normalization, using individual CSF and WM masks obtained from the standard segmentation procedure in SPM12.

### Statistical analysis: fMRI data

Functional images were submitted to a two-stage mixed-effects model (Friston et al., 1994) (GLM). First level analysis was performed using a block design with three regressors of interest (the Relevant and Irrelevant conditions and the rest period) modeled with a boxcar function convolved with the canonical haemodynamic response function. The six motion parameters derived from realignment, the CSF and WM signals and a constant term were included in the model as nuisance regressors. A high-pass filter with cutoff of 512 seconds was applied to the time-series of functional images in order to remove low-frequency noise while preserving signals at task-related frequencies.

Task-related activation maps were obtained from two contrasts of interest. In the Task vs. Rest contrast, we computed the difference between activity during the two conditions (Relevant and Irrelevant) and the rest period. In the “Irrelevant vs. Relevant” contrast, we computed the difference between conditions. The combination of these contrasts allowed identifying functional areas that were engaged by the two tasks but also showing differential activity across conditions. The resulting statistical maps were submitted to second-level group analysis consisting of voxel-wise comparison across subject (one-sample *t*-test), treating each subject as a random effect. Statistical significance was assessed at the group-level using an uncorrected voxel-based threshold of *p* < 0.0001 and a minimum cluster size of 5 voxels. From the resulting group-statistic we selected twenty functional peaks of interest (fourteen from the Task vs. Rest, and six from the Irrelevant vs. Relevant contrast; see Table 1) corresponding to voxels of maximum T-statistic within spatially segregated clusters and uniquely labeled regions (based on the AAL2 and Neuromorphometrics labels, SPM12). Only cortical and sub-cortical regions were considered. In order to extract EEG source activity (see below) from the twenty nodes of interest, all peaks were unwarped back from standard to individual space by applying the deformation parameters generated after normalization.

**Table 1.**
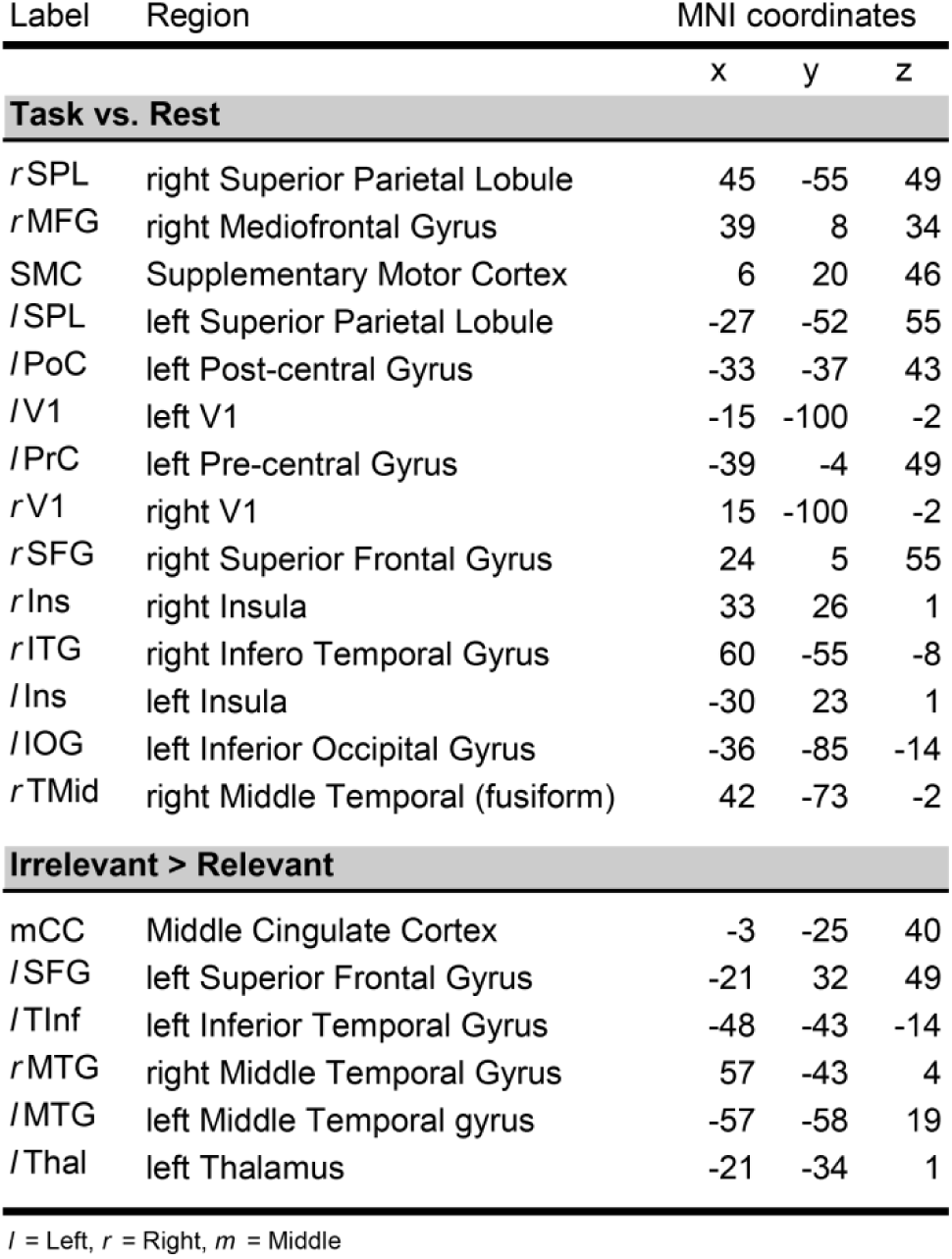
Maxima of fMRI activation from the group statistic on the contrast Task vs. Rest and Irrelevant > Relevant. Only cortical and subcortical peaks within spatially segregated clusters and uniquely labeled regions were selected and used to define ROIs for connectivity analysis. Regions from the Relevant > Irrelevant contrast were not included because of their proximity and overlap with ROIs from the other two contrasts. Regions are labeled using the Neuromorphometrics labeling in SPM12.

### EEG data acquisition and preprocessing

EEG data were acquired on a separate day with a 256 channel EGI Geodesic setup (EGI Eugene, OR, USA). Recordings were digitized at 1000 Hz and referenced against the Cz electrode, the impedance was kept below 50 kΩ. Electrode positions were digitized in 3D using a photogrammetry system (EGI Eugene, OR, USA). The cheek and lower neck electrodes were excluded, leaving 204 electrodes for further analysis. Four participants were excluded from further analysis due to excessive noise in the data. For the remaining twelve participants, EEG recordings were preprocessed using a combination of functions from EEGLAB (Delorme & Makeig, 2004) and custom scripts in Matlab (The Mathworks, Natick, MA).

Prior to signal preprocessing, noisy EEG channels (as determined through careful visual inspection) were removed from the dataset (mean number of channels removed across participants: 19.5±4.6). Data were then down-sampled to 200 Hz, DC-corrected and high-pass filtered at 0.1 Hz with a forward and reverse non-causal FIR filter. EEG epochs were extracted from the continuous dataset and time-locked from -500 ms to 1000 ms relative to the onset of each Gabor. Individual epochs containing non-stereotyped artifacts, peri-stimulus eye blinks and eye movements were identified by visual inspection and removed from further analysis (mean number of epochs removed across participants: 33.4±14.3). Data were then re-referenced to the average potential of all electrodes at each time point and cleaned from line and monitor noise (50 and 75 Hz, plus harmonics) with an adaptive filter (Cleanline plugin for EEGLAB). Remaining physiological artifacts (eye blinks, horizontal and vertical eye movements, muscle potentials and other artifacts) were manually removed through the extended Infomax ICA process implemented in EEGLAB (average number of components obtained across participants: 183.5±4.6; average number of rejected components: 20.3±12.5). After ICA cleaning, excluded channels were interpolated using the nearest-neighbor spline method.

Subject-specific lead fields were computed from a simplified realistic head model (Locally Spherical Model with Anatomical Constraints, LSMAC) derived from individual MRI images, while confining the solution space to the grey matter without constraining source orientation (Brunet, Murray, & Michel, 2011). For each participant, we co-registered the digitized 3D electrode layout with the structural MRI, defining a solution space of about 5000 solution points. Distributed source activity was estimated at each solution point with a linear inverse solution (weighted minimum norm, regularization parameter: 12; Cartool software (Brunet et al., 2011)). Scalar current density values were obtained by projecting instantaneous 3D dipoles to the predominant evoked dipole direction (between 50 and 500 ms after stimulus onset (Coito et al., 2015; Plomp, Leeuwen, & Ioannides, 2010). A single solution point for each functional location of interest was then selected as the one with the higher relative amplitude, determined as the average difference between the magnitude of evoked (between 70 and 150 ms) and baseline (between -100 and 0 ms) activity across trials. Data in the resulting set of twenty solution points of interest were then z-scored across channels and time within participants, and submitted to directed connectivity analysis.

### Statistical analysis: EEG data

The analysis of EEG scalp potentials was restricted to a cluster of electrodes in right and left occipito-parietal regions (n = 110, see Figure 2.A, right panel). This cluster was determined according to previous work showing EEG components related to selective attention at occipital and parietal electrodes, in a comparable experimental design (Anllo-Vento, Luck, & Hillyard, 1998; Daffner et al., 2012). Event-related potentials (ERPs) within the cluster were separately averaged for epochs of the Relevant and Irrelevant conditions. A difference wave was then computed for each participant by subtracting ERPs to the Irrelevant condition from those to the Relevant condition. Individual difference waves were submitted to group-level statistic by means of multiple *t*-tests, comparing the difference at each time point against zero (*p* < 0.05, corrected using false discovery rate).

The power spectral density (see Figure 2.B) was estimated separately for each channel and condition by means of Fourier transformation.

### Directed connectivity

To measure dynamics of directed connectivity between nodes in the Irrelevant and Relevant conditions we used a formulation of partial directed coherence (Baccalá & Sameshima, 2001) (PDC) based on time-varying multivariate autoregressive modeling (tvMVAR) through Kalman filtering (Milde et al., 2010). PDC is a frequency-domain descriptor of directed linear relationships among time series in a network of interacting structures. It is based on the concept of Granger causality (Bressler & Seth, 2011; Granger, 1969) which infers causality in terms of cross-prediction between pairs of signals: signal A Granger-causes signal B if past values of A can be used to improve predictions of future values of B.

The computation of PDC follows the MVAR modeling of a multivariate time series, *Y*, and the estimation of a set of coefficients, *A_(k)_*, describing the linear prediction effect of *k*th past samples of *Y*, *Y_(t-k)_*, on predicting *Y_(t)_*. The *ij^th^* element of *A_(k)_*, *a_ij(k)_*, quantifies directed interactions in the time domain from element *i* to *j*, up to some lag *k*, with *k* > 0 (i.e. excluding instantaneous effects). *A_(k)_* parameters were estimated through a Kalman filter based MVAR approach (Milde et al., 2010), which yields high accuracy in modeling non-stationary, high-dimensional multivariate processes, resulting in estimates of the MVAR model at each time point (tvMVAR).

tvMVAR modeling requires the choice of an optimal lag order *k* and an adaptation constant *c* regulating the trade-off between the speed and stability of tvMVAR estimates (Milde et al., 2010). To determine the optimal combination of *k* and *c*, we used a two-step procedure. In a first step, multiple tvMVAR models of increasing order (from *k* = 1 to *k* = 30) were fitted to all the epochs of each participant using the method of ordinary least squares (OLS). We then used a combination of three information criteria, the Bayesian Information Criterion (BIC), the Akaike Information Criterion (AIC) and the minimum description length (MDL) to estimate the lag order *k* that minimizes each criterion for each participant. A common model order was chosen as the optimal *k* corresponding to the 70^th^ percentile among all *k*s selected by the information criteria, across participants. According to this procedure, we selected a fixed model order of *k* = 10 (50 ms) as a trade-off between sufficient time-frequency resolution and over-parametrization.

In a second step, for each epoch and participant in the two conditions, we computed a series of Kalman filter-based tvMVAR models with fixed model order (*k* = 10) but increasing *c* (from 0.001 to 0.4, in ten logarithmic steps). The model performance at each *c* was evaluated based on two indexes of model fit. The first index is a measure of the goodness-of-fit, the relative explained variance (RExV) (Schlogl, Roberts, & Pfurtscheller, 2000):

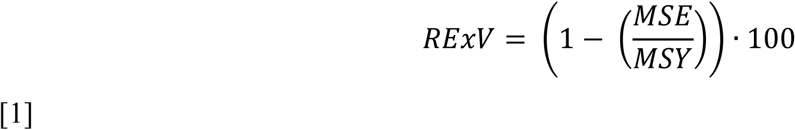

where *MSE* is the mean square of the model residuals and *MSY* is the variance of the multivariate time series *Y*. The RExV indicates the percentage of variance in the signal that is accounted for by the tvMVAR model. The second index, called percent consistency (PC (Ding, Bressler, Yang, & Liang, 2000), measures the percentage of the correlational structure in the data that is captured by the MVAR model and is expressed as:

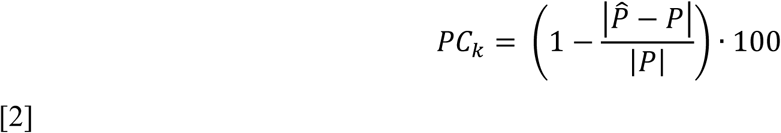

where 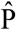 is the vector of all pairwise cross-correlations among signals predicted by the tvMVAR model, up to a lag *k* = 50, and P is the vector of all pairwise cross-correlations in the real data. The RExV and PC for each adaptation constant were averaged across participants and the optimal *c* was selected as the one for which both the RExV and PC where higher than 85%. According to this procedure, we selected an adaptation constant of *c* = 0.1 which is consistent with the range reported in previous work (Astolfi et al., 2008).

Thus, combined together, the model order selection based on information criteria and the choice of *c* based on RExV and PC provided a parsimonious MVAR model to best explain variability and cross-correlational structures in the data. For each participant, the MVAR coefficients and PDC were estimated on individual epochs and epochs for which the MVAR model had PC lower than 70% were discarded from further analysis (mean across participants: 3.9±4.7%).

Time-varying PDC was then computed from the frequency-transformed *A_(k)_* parameters as:

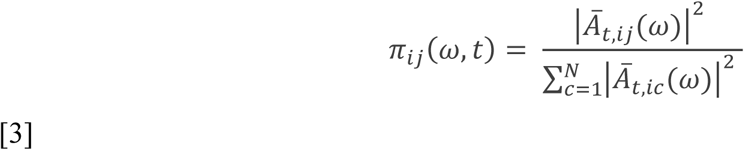

where 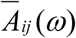 is the spectral representation of the coefficient matrix *A_(k)_*, and *π* quantifies the interaction from element *i* to *j* at frequency *ω* and time *t*. The square exponents enhance the accuracy and stability of the estimates (Astolfi et al., 2006) and the denominator allows the normalization of outgoing connections by the inflows (row-normalization by the sum of each columns *c* of the *i*-th row), which improves the accuracy and physiological plausibility of the results (Plomp, Quairiaux, Michel, & Astolfi, 2014).

As a final step, single-trial, scaled PDC estimates (from 0 to 1) were multiplied by the spectral power (SP) of each node. For each trial and node, SP was computed in the 6-80Hz frequency range using a multitaper Hanning window function with a sliding window of 5 ms and a window length of 250 ms. The obtained SP were scaled (from 0 to 1) and multiplied by PDC at the single epoch level. This yielded a weighted version of PDC (*wPDC*) which embeds information about the predominant frequencies of interaction and whose physiological plausibility has been previously validated in animal models (Plomp et al., 2014, p. 2014).

Single-trial estimates of *wPDC* were then averaged separately across conditions resulting in a set of individual matrices (node x node x time x frequency) summarizing the time-varying directed interactions among all nodes in the Relevant and Irrelevant conditions.

### Statistical analysis: wPDC

Statistical analysis of *wPDC* as a function of the Relevant and Irrelevant conditions was restricted to a time window of interest (from -50 ms to 500 ms from stimulus onset, discarding a number of time frames equivalent to the model order) and performed by means of a non-parametric, cluster-based permutation test which controlled over the false positive rate in a multiple comparison setting (Maris & Oostenveld, 2007). The general approach is as follows: for the contrast Relevant minus Irrelevant, a paired *t*-test across participants is computed for each time-frequency point of each cell of the *wPDC* matrix. The resulting statistic is then thresholded (here at *p* < 0.05) and adjacent significant time-frequency points are combined to form clusters. For each cluster, a new statistic is then obtained as the sum of all *T*-values in the time-frequency points that define the cluster. This procedure was repeated (n = 5000) while shuffling conditions across participants and retaining the maximum cluster-forming *T*-value obtained. The proportion of surrogate clusters with maximum *T* larger than the one observed defines the corrected *p* value (maximum *T*-statistic). The permutation test was performed to identify clusters of significant differences in *wPDC* between conditions (*p* < 0.05) in the 6-80 Hz frequency range.

We summarized results from the thresholded wPDC matrix as follows. First, we separately summed all significant interactions across nodes in the Relevant and Irrelevant condition and we combined them into a single matrix, summarizing global network dynamics and frequency distribution as a function of whether the Gabor stimulus was attended or ignored (Figure 3.A). As a second step, we computed the summed outflow for each node in the condition showing the larger number of significant interactions (the Irrelevant condition). The summed outflow was calculated as the sum of significant outgoing wPDC values from each node at each time point, averaged across a frequency band of interest (α-band, 6±16 Hz, see Figure 3.B). In a final step, we assessed the direction of information flow across time from the two predominant drivers of α activity in the Irrelevant condition (left and right SPL, see Figure 3.C). Three directions of interest were defined according to the relative position in the axial plane of each receiver node with respect to the two parietal drivers: an “interhemispheric” direction, assessing the degree of cross-interaction between the two nodes of interest, a “feedback” and a “feedforward” direction, summarizing the amount of parietal driving toward posterior and anterior nodes, respectively. Only cortical nodes were included in this summary.

### Phase-amplitude coupling

Several algorithms have been proposed to extract phase-amplitude coupling measures based on different approaches (Dvorak & Fenton, 2014b). Here we used the raw modulation index (*MI_raw_*) (Canolty & Knight, 2010; Onslow, Bogacz, & Jones, 2011; Penny, Duzel, Miller, & Ojemann, 2008) which has better statistical properties than the normalized MI (Penny et al., 2008) and because within-condition normalization or surrogate statistics is not suitable for comparing PAC between experimental conditions. The *MI_raw_* was computed for two nodes of interest (left and right V1). Source activity in the two nodes was filtered at a set of frequencies in the α (6-16 Hz, in steps of 1 Hz) and γ-band (50-80 Hz, in steps of 2 Hz) via convolution with complex Morlet Wavelets (width = 7) (Lachaux, Rodriguez, Martinerie, Varela, & others, 1999; Onslow et al., 2011). To avoid edge artifacts, filtering was applied on a time window of interest (from 0 to 500 ms after stimulus onset) plus additional buffer windows of 500 ms before and after. The instantaneous low-frequency phase (*Φ_α_*) and high-frequency amplitude (*A_γ_*) were extracted from the filtered waves and a composite signal

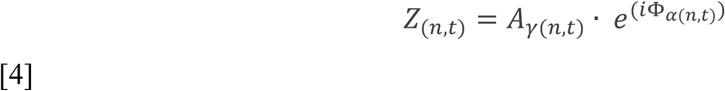

was created at each time point *t* and epoch *n*, for all combinations of low and high frequencies of interest. The *MI_raw_* index was then calculated separately for the two conditions as

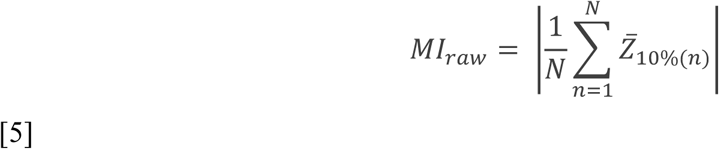

where *n*[1, …, *N*] are trials of each condition and 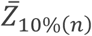 denotes the 10% trimmed mean length of vector *Z*, across time bins *t*, which we found to be more robust and less sensitive to spurious coupling. This procedure yielded a set of individual matrices quantifying the degree of phase-amplitude coupling among α and γ frequencies in the two conditions. Statistical testing of the difference between conditions (Relevant minus Irrelevant, see Figure 4.B, left panel) was performed at the group-level using permutations (n = 50000) and the maximum *T*-statistic applied over the whole matrix.

To generate Figure 4.B (right panel), we computed the phase-locking value (*PLV*) (Lachaux et al., 1999; Penny et al., 2008) between all frequencies in the γ-band and the subset of frequencies in the α-band showing significant differences in the *MI_raw_* across conditions. The PLV was computed for each condition at each time bin *t* as:

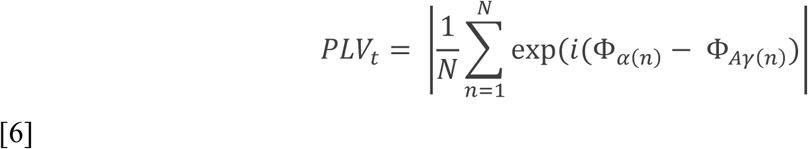

where Φ_*Aγ*(*n*)_ is the instantaneous phase of the amplitude-filtered high frequency signal in trial *n*. The difference in *PLV* across conditions (Δ*PLV*) was calculated by subtracting values of the Irrelevant condition to the Relevant condition at each time point. An exploratory analysis was performed on Δ*PLV* by means of permutation test (*p* (uncorrected) < 0.05, two-tailed), shuffling the labels of conditions across participants 50000 times.

To test whether local changes in phase-amplitude coupling at occipital nodes were related to the incoming parietal drive in the α-band (see Results section), we ran a mixed-effects linear regression analysis using the *fitlme* procedure with maximum likelihood estimation in Matlab (Statistics and Machine Learning Toolbox). Mixed-effects linear regression analysis allows the mixing of categorical and continuous variables and provides a better account of inter-subject variability than canonical repeated-measures ANOVA models. Log-transformed, single-trial estimates of *MI_raw_* (the absolute of 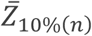 at frequency tiles of significant differences, *Φ_α_* = 10-11 Hz; *A_γ_* = 58-64 Hz) were modeled with the continuous predictor Parietal α*_drive_* (the single-trial sum of incoming α drive from parietal nodes in the time window between 0 and 500 ms post-stimulus), the factor Condition (Relevant vs. Irrelevant) and their interaction (Condition* Parietal α*_drive_*), incorporating subjects as random effect over the intercept term. With a similar approach, we also tested for linear relationships between the amount of PAC at occipital nodes and the outgoing drive in the γ-band from these regions. In this second model, occipital single-trial estimates of *MI_raw_*, the factor Condition (Relevant vs. Irrelevant) and their interaction (Condition\**MI_raw_*) were used to predict the log-transformed sum of occipital drive in the γ-band (Occipital γ_*drive*_, 50-80 Hz).

## Results

### Visually evoked potentials and power spectrum

The behavioral task is depicted in Figure 1. Participants fixated on a central spot while presented with sequences of oriented Gabors. They were instructed either to attend to each Gabor, reporting its orientation (Relevant condition), or to ignore the Gabors and report an occasional change of color in the fixation spot (Irrelevant condition). Thus, participants were exposed to near-identical visual stimulation across conditions, but under different task demands.

**Figure 1.**
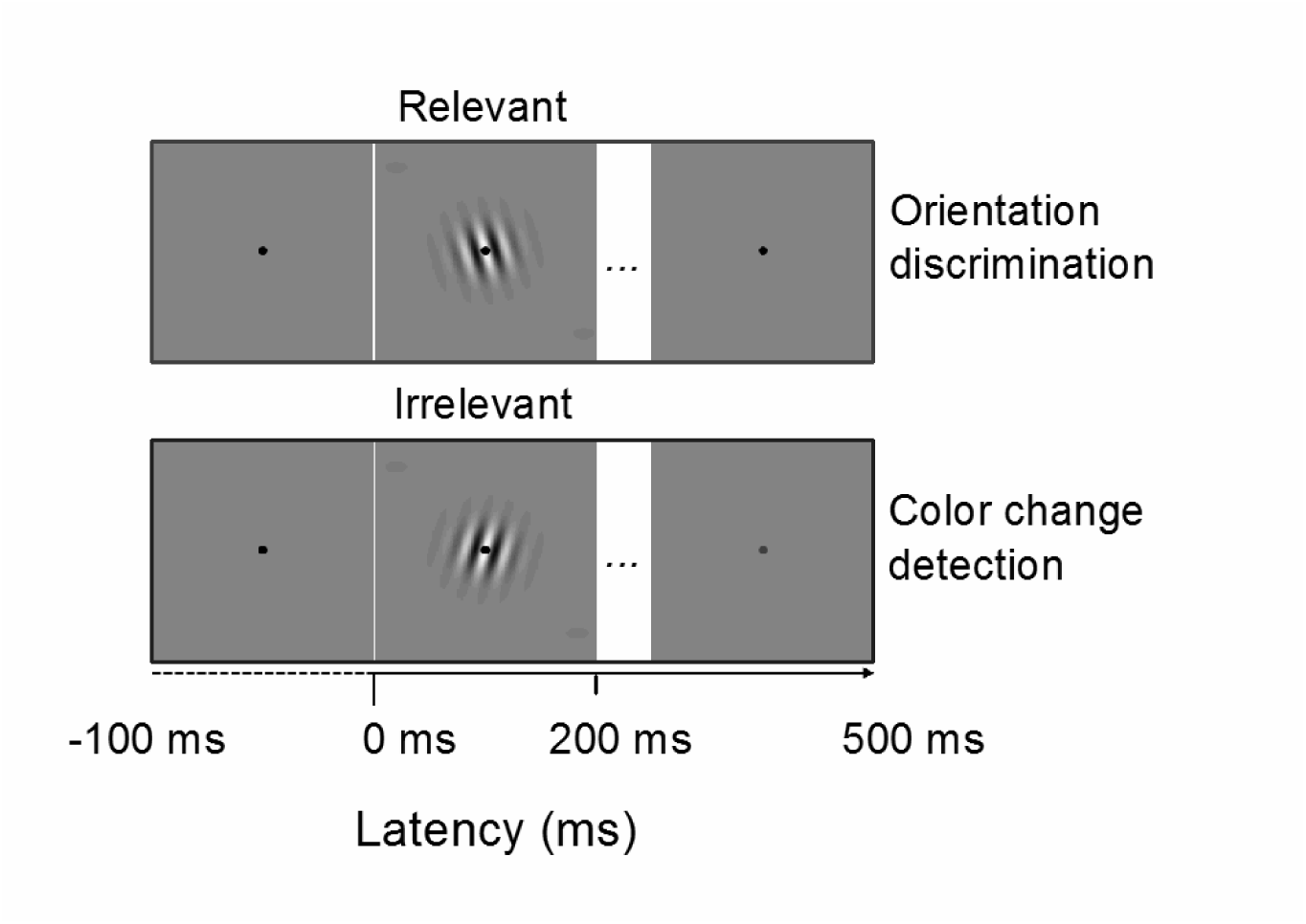
Experimental paradigm. In the Relevant condition, participants discriminate the orientation (left vs. right) of briefly presented Gabor stimuli. In the Irrelevant condition, Gabor stimuli were ignored and participants had to report an occasional color change in the fixation spot. Epochs for the EEG and connectivity analysis were time-locked to the onset of each Gabor. Stimuli are not drawn to scale.

We first investigated the effect of selective attention at the EEG electrode level. Based on previous reports using similar attentional manipulations, we expected a SN component, which has been extensively related to attentional selection (Daffner et al., 2012; Hillyard & Anllo-Vento, 1998). The SN is observed in difference potentials (i.e., by subtracting the unattended to the attended condition) at posterior-occipital electrodes. Thus, we restricted our analysis to a set of posterior-occipital electrodes (see Figure 2.A, right panel) and we subtracted the averaged evoked responses across these electrodes in the Irrelevant condition from those in the Relevant condition. Figure 2.A shows a first significant positive difference across conditions at the time of early visual evoked responses (from 95 to 105 ms post-stimulus, *p*(FDR) < 0.05) followed by the expected SN component, emerging at 225 ms and persisting for the rest of the epoch (first significant time window from 225 to 275 ms, second time window from 315 to 430 ms and third time window from 460 to 495 ms), consistent with previous reports (Daffner et al., 2012; Hillyard & Anllo-Vento, 1998). For Relevant stimuli the grand average α-band power was reduced (6 to 12 Hz, peak at 10 Hz, see Figure 2.B), with largest reductions at parieto-occipital electrodes. This shows that our attentional manipulation was successful in eliciting an SN.

**Figure 2.**
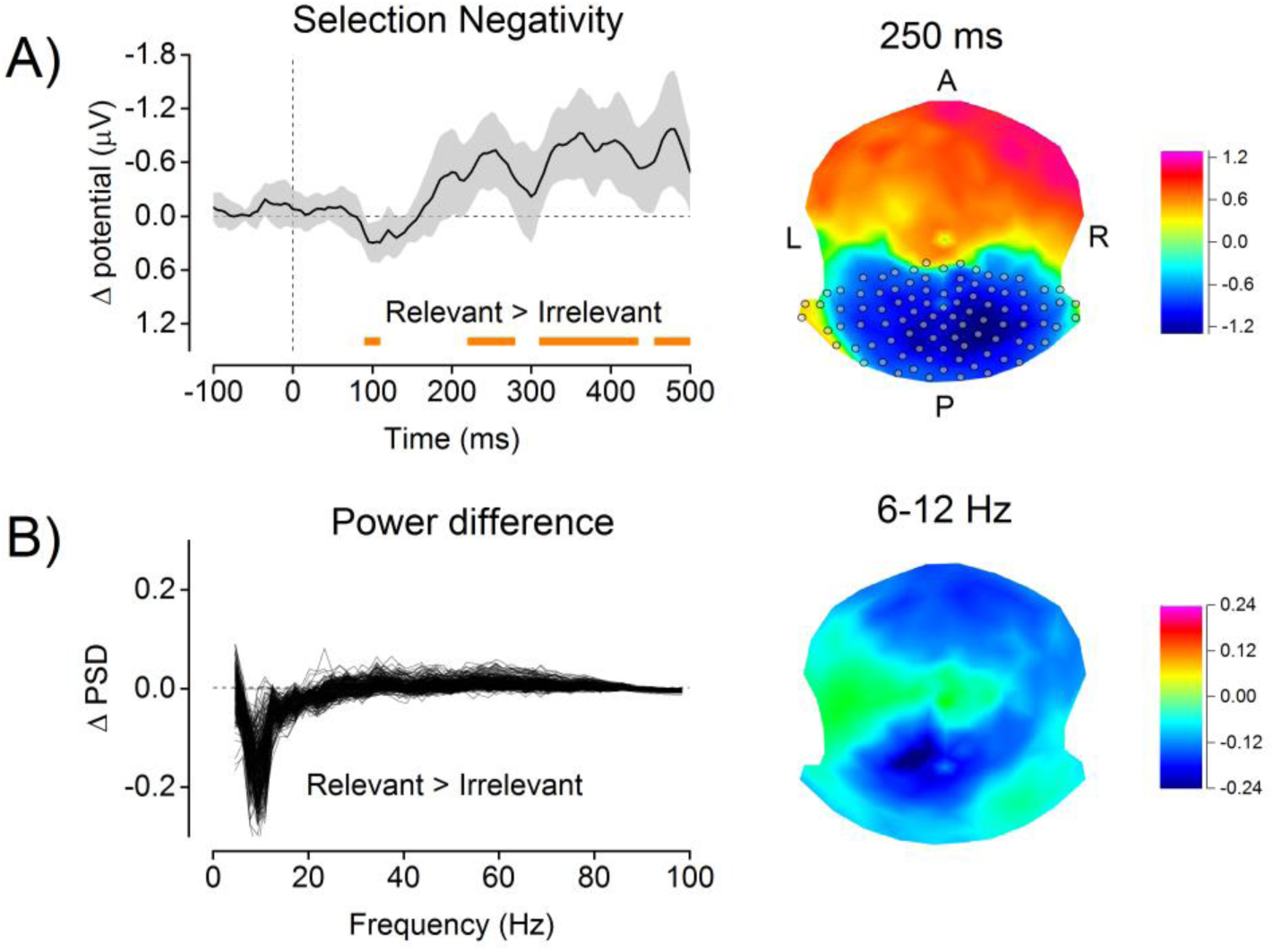
Attentional modulations of evoked potentials and power spectrum. A) The Selection Negativity (SN) component (left panel) observed in a subset of occipito-parietal electrodes (gray circles, right panel) and the corresponding SN scalp topography at 250 ms (right panel). Significant differences between evoked activity in the Relevant and Irrelevant condition are highlighted by the orange bottom line (right panel, *p*(FDR) = 0.05). Gray shaded areas are 95% confidence intervals of the mean. B) The difference in the overall power spectrum between conditions for all EEG channels (Relevant minus Irrelevant, left panel) and the topography of the increased α activity (6-12 Hz) in the Irrelevant condition at posterior sites (right panel).

### Source-level connectivity

To investigate brain dynamics of directed interactions as a function of selective attention we first identified twenty nodes from functional imaging (Methods, Supplementary figure 1). This resulted in a set of nodes (Table 1) including primary visual cortex, lateral occipital and middle temporal cortex, parietal and frontal cortex, insular regions and cingular cortex, in line with perceptual and attentional networks characterized by previous work (Corbetta & Shulman, 2002).

The results of the source-connectivity analysis are summarized in Figure 3.A. This figure shows the sum of the connectivity strengths across all nodes that significantly varied with selective attention, in the time and frequency domains. On average, increased connectivity for Irrelevant stimuli occurred in the α-band (6 to 16 Hz), whereas for relevant stimuli connectivity increased in the γ-band (50 to 80 Hz). Network dynamics in the γ-band emerged rapidly after stimulus onset, at latencies of early visual evoked responses (first peak at ~60 ms after stimulus onset) and showed a maximum at 220 ms, followed by a gradual decrease. In the Irrelevant condition, significant directed interactions in the α-band occurred at longer latencies (from ˜ 100 ms post-stimulus) and increased during the rest of the epoch.

**Figure 3.**
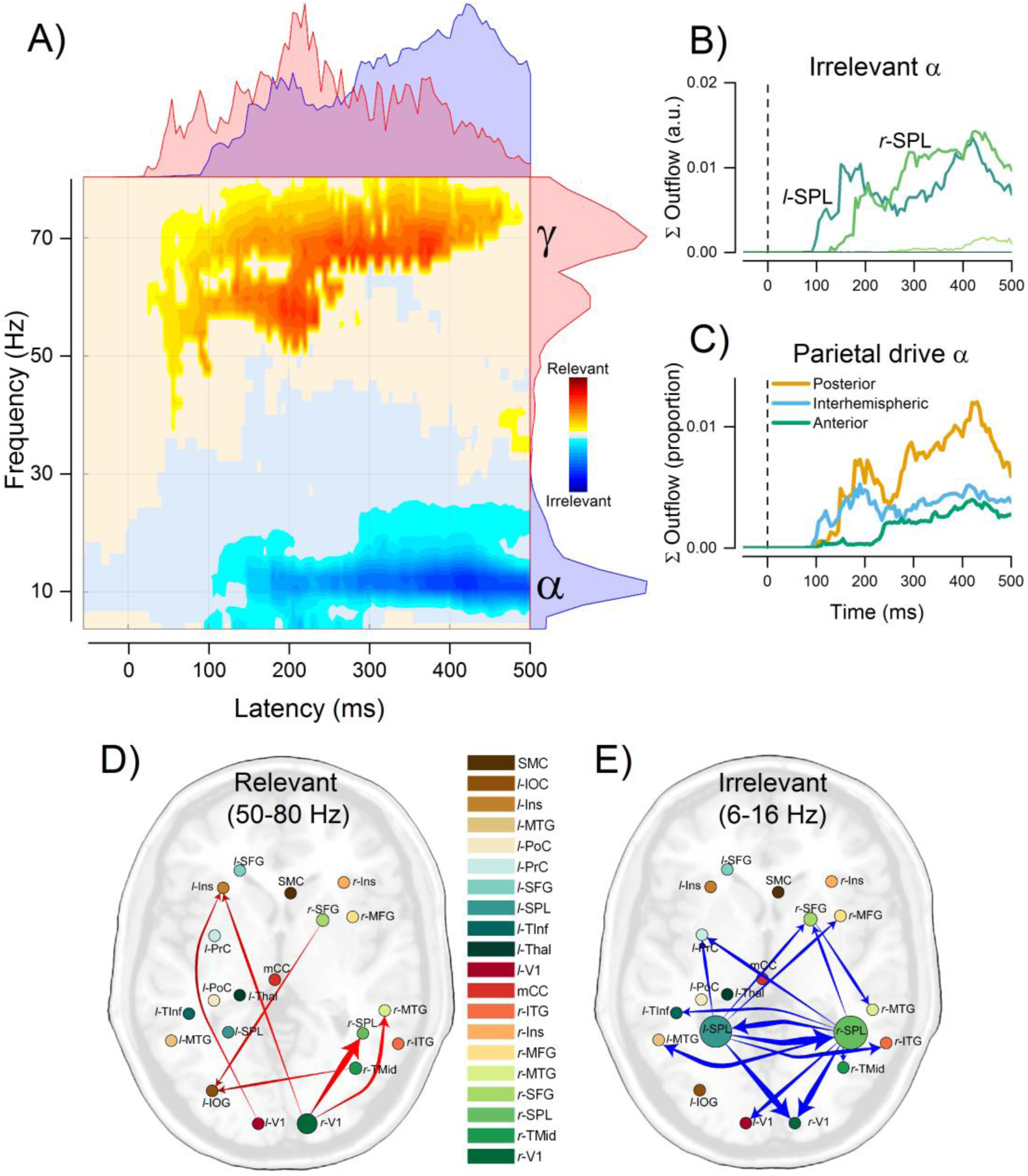
Time-varying directed connectivity results. A) Time-frequency plot summarizing the difference in global network dynamics across conditions. The summed strength of all interactions that showed significant attentional modulation (*p* < 0.05, cluster-based correction for multiple comparisons) is shown in the yellow-red color scale for the Relevant > Irrelevant comparison and in the cyan-blue color scale for the Irrelevant > Relevant comparison. Marginal plots on the right represent time-collapsed frequency distributions of significant interactions. Marginal plots on the top are frequency-collapsed distributions of interactions over time. Marginal distributions are plotted separately for the Relevant (red) and Irrelevant (blue) condition. B) Sum of the total α outflow (6-16 Hz) in time from each node in the Irrelevant condition. C) Direction of α interactions from parietal nodes over time. D) Average of significantly increased interactions in the γ-band during the Relevant condition. E) α-band connections with average strength below 1% of the total network are not shown.

Relevant stimuli evoked directed interactions in the γ-band from early visual nodes to regions in the lateral-occipital cortex, middle-temporal cortex, right superior parietal cortex, and from a frontal node to the occipital cortex (Figure 3.D). The significantly increased high-frequency interactions in the Relevant condition almost exclusively originated from early visual areas (rV1(95.82%), rTMid(2.74%), lV1(0.06%)). A more complex pattern of significant connections was found in the Irrelevant condition with an elevated number of reciprocal interactions (see Figure 3.E). Figure 3.B represents the sum of outflow across significant time-frequency points in the α-band, for each sender node. This summary identified the left and right parietal cortex as the main drivers in the α-band. Significantly increased α-band driving from parietal cortex emerged after about 100 ms from stimulus onset, with initial peaks between 100 and 200 ms and a later peak after 400 ms, targeting a wide range of nodes.

To further characterize the dynamics of parietal driving in the Irrelevant condition, we divided receiving nodes according to their relative position in the axial plane, with respect to the parietal senders. Three main directions were defined: posterior (feedback), anterior (feedforward) and interhemispheric (between the parietal nodes), The dynamics summed outflow from parietal nodes in these three directions is summarized in Figure 3.C. Parietal interactions were characterized by an initial interhemispheric coupling between the two nodes, emerging at around 100 ms from stimulus onset, followed by a predominant and gradually increasing drive to posterior regions, and a late increase in driving toward anterior regions. A consistent portion of the outgoing α drive from parietal nodes was interhemispheric or directed toward visual processing areas (interhemispheric (29.89%), rV1 (29.03%), lMTG (8.68%), rITG (7.06%), lV1 (6.07%)). During the initial interhemispheric drive (100-200 ms), 60% of significant bi-directional interactions were directed from the left PPC (*l*-PPC) to the right (*r*-PPC) and 40% in the opposite direction.

### Occipital phase amplitude coupling

The connectivity analysis revealed two main findings: 1) Selectively attending to stimuli increases the outflow of γ activity from early visual areas and 2) ignoring stimuli evokes distributed interactions in the α-band, orchestrated by superior parietal areas and directed predominantly toward nodes in the primary visual cortex. Following these results, we investigated whether oscillatory influences from parietal cortex interact with local mechanisms that coordinate the routing of information carried by high-frequency oscillations from visual areas. We tested this hypothesis by investigating the impact of parietal α drive on the degree of local PAC at early visual nodes. PAC reflects dependencies between the phase of a low-frequency oscillation and the amplitude of the high-frequency component of a neural signal (Szczepanski et al., 2014), and has been proposed as a mechanism of gating (Bonnefond & Jensen, 2015; Jensen & Mazaheri, 2010) and local coordination of neural processing (Dvorak & Fenton, 2014a), by which low frequencies modulate the excitability of neuronal ensembles (Canolty & Knight, 2010).

In line with this idea, we hypothesized that α-γ coupling in visual regions would be higher for Relevant stimuli, reflecting increased processing and information flow from these nodes. To test this hypothesis, we compared PAC across conditions (see Methods) for each frequency point in the range of α -and γ-bands, as identified from the connectivity analysis. As expected, we found an overall tendency for stronger α-γ coupling in the Relevant than in the Irrelevant condition with a significant difference in the PAC between phases at 10-11 Hz and amplitudes at 58-64 Hz (*p* < 0.05, two-tailed t-test, see Figure 4.B, left panel). As shown by the phase locking value (PLV, see Methods) between phases at 10-11 Hz and γ amplitudes (58-64 Hz), this increased coupling emerged between 100 ms and 200 ms from stimulus onset (see Figure 4.B, right panel), consistent with the first window of attentional modulation of connectivity, and occurred without statistically significant differences in α (*p* = 0.206, two-tailed t-test) and γ power (*p* = 0.203) between conditions.

**Figure 4.**
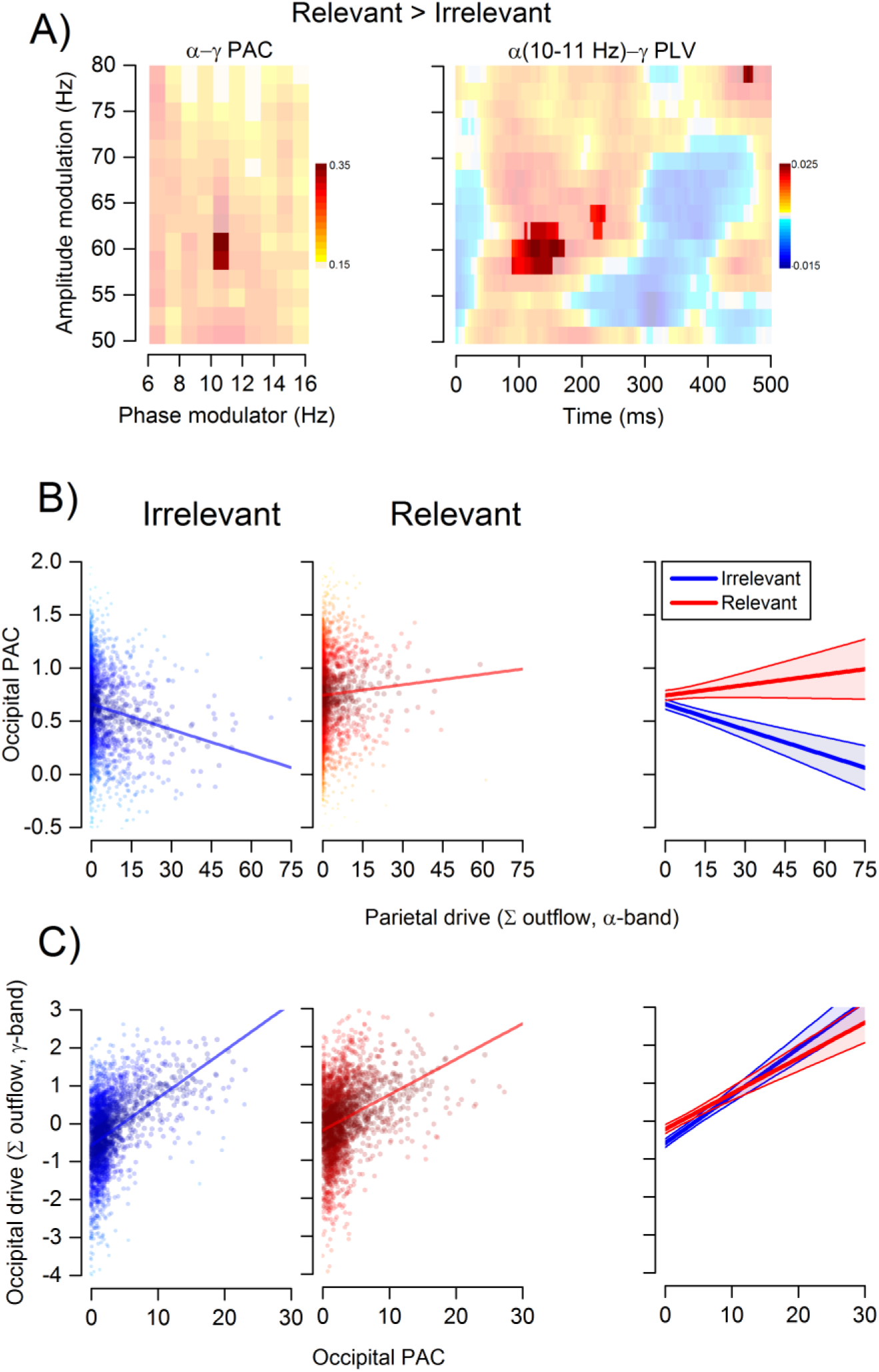
Parieto-occipital interactions affect occipital phase amplitude coupling. A) Differences in occipital PAC (left panel) and PLV (right panel) across conditions. PAC is shown for low-frequency phases (6-16 Hz) and high-frequency amplitudes (50-80 Hz) of interest. PLV is shown for low-frequencies phases of significant PAC (10-11 Hz) and high-frequencies amplitudes (50-80 Hz). Opaque areas represent regions of significant PAC (*p* < 0.05, maximum T-statistic) and PLV (*p*(unc.) < 0.05, permutation-based statistic). B) Relationship between the summed parietal α outflow to V1 and the PAC measured at occipital sites for the Irrelevant (left panel, blue dots and line) and Relevant (middle panel, red dots and line) condition. The right panel shows the interaction Condition x Parietal α-drive. C) Relationship between the occipital PAC and the summed γ outflow from occipital nodes for the Irrelevant (left panel) and Relevant (middle panel) condition. The right panel shows the interaction Condition x Occipital PAC. In panel B) and C), dots are single trial values of all participants, with individual random intercepts removed. Lines and shaded areas are the estimated regression lines and 95% prediction intervals from each multilevel model. Darker colors reflect points closer to the predicted regression line.

### PAC mediates γ driving from early visual nodes

Hypothesizing that phase-amplitude coupling mediates the effective outflow of early visual areas, we predicted 1) that the amount of incoming α-drive from parietal nodes should predict the local PAC and 2) that PAC predicts outgoing connection strengths in the γ-band. To test this, we used two multilevel linear models (see Methods). In the first model, we modeled α-γ occipital PAC as a function of Parietal α*_drive_*, the Condition factor (Irrelevant vs. Relevant) and their interaction. The model revealed a significant intercept (β = 0.659±0.260, *p* < 0.05, Satterthwaite approximation for degrees of freedom), a significant negative slope for the continuous predictor Parietal α_*drive*_ (β = -0.007±0.001, *p* < 0.001), a significant main effect of Condition (β = 0.082±0.010, *p* < 0.01) and a significant interaction between Parietal α_*drive*_ and Condition (β = 0.011±0.002, *p* < 0.001). The results of this multilevel analysis showed 1) a decrease of occipital PAC with the increasing parietal drive during Irrelevant trials, 2) an overall higher PAC in the Relevant condition and 3) that Parietal α_*drive*_ markedly decreased occipital PAC for Irrelevant trials while it increased occipital PAC on Relevant trials (see Figure 4.C).

Following a similar approach, in a second model we tested the hypothesis that the amount of α-γ PAC at visual nodes could predict the degree of information flow from these regions, as indexed by the sum of occipital outgoing interactions in the γ-band. This second model had a non-significant intercept (β = -0.581±0.641, *p* > 0.05) and a significant positive slope for the continuous predictor occipital PAC (β = 0.124±0.009, *p* < 0.001), indicating that γ interactions from early visual nodes increased as a function of the strength of local PAC. We also obtained a main effect of Condition (β = 0.368±0.040, *p* < 0.001) and a significant interaction between occipital PAC and Condition (β = -0.031±0.008, *p* < 0.001), revealing that, although outflow γ interactions were overall higher in the Relevant conditions, increases in the amount of occipital PAC led to larger γ outflow in the Irrelevant condition (see Figure 4.D). Because confidence intervals of the regression widened considerably with increased PAC values, due to fewer observations, we refrain from interpreting this interaction further.

## Discussion

Our results are compelling evidence that selective attention emerges from the interplay between frequency-specific, large-scale interactions and local dynamics of neuronal computation. Our main findings can be summarized as follows: 1) selective attention increases early influences from visual nodes to higher-level areas in the γ-band; 2) suppression of task-irrelevant stimuli is orchestrated by parietal nodes through top-down driving in the α-band; 3) influences from parietal nodes determine nested oscillations and signal outflow from occipital sites by changing local α-γ couplings.

The finding of predominant γ interactions in response to attended stimuli is well in agreement with the current understanding of γ activity as a marker of neuronal excitability and functional integration in large-scale networks dedicated to perceptual processing (Bastos et al., 2015; Fries et al., 2001; Gruber et al., 1999; Jensen et al., 2007; Michalareas et al., 2016; Müller, Gruber, & Keil, 2000; Tallon-Baudry et al., 2004; Vossel et al., 2014). Post-stimulus γ oscillations in visual cortex have been shown to correlate with neuronal spikes (Belitski et al., 2008) and to foster neuronal gain mechanisms and behavioural performance (Ni et al., 2016; Womelsdorf, Fries, Mitra, & Desimone, 2006). Furthermore, recent work in primates and humans has shown that influences along feedforward projections are dominated by γ-band synchronization, suggesting a pivotal role for high frequencies in the propagation of task-relevant signals (Bastos et al., 2015; Buffalo et al., 2011; Michalareas et al., 2016; Van Kerkoerle et al., 2014). Here we provide the first dynamic, high-resolution temporal investigation showing that task-relevant processing evokes directed influences in the γ-band from early visual areas at very short latencies. These rapid interactions emerged with initial peaks at latencies of visual evoked responses (~90 ms) and propagated in the feedforward direction, targeting regions involved in higher level visual processing (i.e., lateral occipital cortex and middle temporal gyrus (Grill-Spector, Kourtzi, & Kanwisher, 2001; Ungerleider & Haxby, 1994)) visual attention (right parietal lobule (Corbetta & Shulman, 2002)) and response selection/evaluation (left insular cortex (Corbetta, Miezin, Dobmeyer, Shulman, & Petersen, 1991; Eckert et al., 2009; Menon & Uddin, 2010)). By characterizing the fast dynamics with which γ oscillations route attended information along the ascending pathway, we expand on previous work relating attention to increased γ activity (Müller et al., 2000) and synchronization (Fries et al., 2001; Fries, Womelsdorf, Oostenveld, & Desimone, 2008).

Whereas γ-band interactions increased with attentive stimulus processing, irrelevant stimuli triggered wide-spread influences in the α-band. Recent insights on the functional role of α activity have suggested its involvement in mediating inhibition of cortical structures activated by irrelevant events (Herrmann, Strüber, Helfrich, & Engel, 2016; Jensen & Mazaheri, 2010; Klimesch, 2012; Rihs, Michel, & Thut, 2007). Contrary to high frequencies, α oscillations are inversely related to attention and behavioural performance, their amplitude increases in regions associated with irrelevant processes and decreases in areas engaged by relevant and attended stimuli (Jensen & Mazaheri, 2010; Klimesch, 2012). Therefore, α-band synchronization has been proposed as a mechanism to prevent local neural analyses and network propagation of stimulus information (Womelsdorf & Fries, 2008). Here we found distributed interactions in the α band that began at around 100 ms after stimulus onset and persisted until the end of each epoch (500 ms), thus encompassing early and later stages of perceptual processing. This result is consistent with the emergence of a stimulus-evoked brain-wide inhibitory drive through α oscillations that prevents multiple brain regions from receiving irrelevant and interfering information.

It is interesting to note that the time course of α interactions roughly paralleled the selection negativity (Hillyard & Anllo-Vento, 1998) component observed at the sensor level. Our findings, therefore, suggest that the well-known SN might result from increased parietal driving that inhibits the information transfer from visual areas.

As for the sources of such top-down inhibitory drive, our study is the first to reveal the key driving role of the posterior parietal cortex (PPC) from EEG recordings. This result is in line with the established role of PPC in coordinating shifts of spatial attention and disengagements of attention from stimuli (Posner & Petersen, 1990) and with more recent studies suggesting its top-down regulatory effect on visual selection (Bressler, Tang, Sylvester, Shulman, & Corbetta, 2008; Hung, Driver, & Walsh, 2005; Kastner & Ungerleider, 2001; Plomp, Hervais-Adelman, Astolfi, & Michel, 2015; Yantis et al., 2002)

We show that α-mediated inhibitory signals originated from sources in the superior portion of the PPC. These suppressive influences emerged swiftly after stimulus onset, first targeting regions of early perceptual processing and then interacting with nodes in the frontal cortex at longer latencies (Figure 3). This suggests that the PPC may have the key function to prevent irrelevant processing in a hierarchical fashion, from the inhibition of early perceptual analyses to the tracing and suppression of residual signals at later stages. We should caution that although we functionally identified these sources in a superior portion of the parietal cortex using fMRI, the low spatial resolution of EEG and potential residual mixing of signals after source reconstruction, prevent the complete separation of activity in these regions from that in neighbouring loci, such as those reported by previous work (Plomp et al., 2015; Posner & Petersen, 1990) and thus, here we broadly refer to the parietal nodes as the PPC.

The first pattern of α-band driving from parietal cortex was interhemispheric, targeting the homologous region in the contralateral hemisphere, with stronger interactions from the left to the right PPC. Selective inhibition of visual information may therefore require initial interhemispheric interactions that set the parietal system into an inhibitory state, after which PPC effectuates attentional suppression is through long-range interactions. This interpretation is consistent with recent work showing that inhibitory interactions induced by bilateral TMS over PPC prevents the parietal system from promoting the excitability of visual areas (Silvanto, Muggleton, Lavie, & Walsh, 2008).

In addition to demonstrating the critical role and the dynamics of α-band interactions from PPC, our results suggest a mechanism for how parietal cortex modulates activity in distal areas during selective attention. We demonstrated, for the first time, a direct link between the amount of incoming α modulation and the degree of phase-amplitude coupling (PAC) in occipital areas. PAC reflects the coupling of the amplitude of high-frequency oscillations to the phase of slower components (Szczepanski et al., 2014) and has been positively related to neuronal processing and communication (Canolty & Knight, 2010; Dvorak & Fenton, 2014a; Voytek et al., 2010). PAC can serve as an information integration mechanism over multiple temporal and spatial scales. During PAC, the amplitude of a fast rhythm (e.g., γ oscillations), and the potentially related degree of neuronal excitability, increases at specific phases of a slow oscillation (e.g., α rhythm). This system of nested oscillations implies that low frequencies rhythmically modulate and coordinate neuronal activity, optimizing information processing within ensembles of neurons (Dvorak & Fenton, 2014a) and favoring the exchange of relevant information among distant ensembles oscillating in synchrony (Voytek et al., 2010).

Our work demonstrates a strong relationship between parietal drive in the α range, local occipital PAC, and γ-band driving. Previous work has emphasized the potential role of PAC as a mechanism for communication within and between cortical areas under attentional demands (Sadeh, Szczepanski, Knight, & Mangun, 2014), suggesting a potential target mechanism for the influence of top-down selective modulation. In line with this view, our findings show that attentive processing of stimuli leads to increased α-γ PAC in visual cortex, which in turn, predicts the degree of occipital outgoing γ interactions. Ignoring a stimulus, on the other hand, leads to decreased occipital α-γ PAC, and this decrease is determined by the incoming parietal inhibitory drive. These results point to an important functional interpretation of the role of PPC in selective attention: to ignore irrelevant information, Parietal inhibitory α interactions disrupt structures of information processing at target regions by decoupling fast from slow oscillations and thus, reducing the influences that these regions exert on upstream areas. This way, local PAC in V1 works as a mechanism for signal propagation to upstream areas, while PPC gates neuronal communication by adjusting occipital PAC as a function of attentional demands through directed interactions in the α-band.

Our findings provide empirical support for a recently proposed account of cross-frequency neuronal communication (Bonnefond, Kastner, & Jensen, 2017). On this theory, neuronal communication occurs through synchronization between areas at low frequencies, which in turn aligns the excitability phase of different pools of local neurons. According to this framework, neuronal ensembles exchanging task-relevant information would oscillate coherently in the α-band with a collective decrease in α power. The low α amplitude would allow longer windows of shared excitability among synchronized regions, favoring interactions in the γ-band. This predicts that attentional suppression may be accompanied by two phenomena: 1) a substantial increase in α power and 2) a loss of α synchrony among local and distal neuronal pools processing task-irrelevant signals, with a consequent reduction of their γ-mediated interactions. Whereas our findings are broadly in line with this perspective, they tend to favor the latter mechanism as the primary mediator of attentional inhibition. It is worth emphasizing that, at the source level, occipital α power was not significantly increased by the occurrence of irrelevant stimuli, nevertheless, γ-band interactions and occipital PAC were both decreased (Dvorak & Fenton, 2014b). This suggests that attentional suppression may be the product of an induced non-rhythmic (e.g., de-synchronized) behavior in the excitability profile of neuronal ensembles and a resulting failure to establish coherent, high-frequency interactions with other regions. Crucially, our results indicate that such de-synchronization of occipital activity may be induced by parietal neurons through fast and directed interactions in the α-band.

Although our paradigm did not allow a direct (e.g., behavioral) measure of participants’ attention, similar protocols have been widely used in the fMRI (Sunaert, Van Hecke, Marchal, & Orban, 2000) and EEG (Daffner et al., 2012; Hillyard & Anllo-Vento, 1998) literature to investigate mechanisms of selective attention underlying the processing of task-relevant or irrelevant stimuli. An interesting question for future studies would be to extend our findings to other classical attentional task (i.e., the Posner paradigm) in which behavioral performance can be directly related to network dynamics at critical frequency bands. In a similar vein, we focused on two specific frequency bands identified in a data-driven way. This does not exclude that in other paradigms, under different attentional manipulations, other oscillatory components may play critical roles (Siegel et al., 2008; Szczepanski et al., 2014).

In conclusion, our findings support the view of selective attention as a top-down modulation of oscillatory activity (Womelsdorf & Fries, 2008) and provide first evidence of the causal role of α-band influences from parietal cortex in biasing rhythmic synchronization and neural efficacy at target regions.

## Data and code availability

The data and code generated during the current study are openly available from the corresponding author on reasonable request.

## Acknowledgements

This work was supported by Swiss National Science Foundation grants to GP (PZ00P3_131731 and PP00P1_157420/1).

